# The efficiency of different transmission routes of *Xanthomonas citri* pv. *fuscans* and other seed-borne bacteria to bean seeds

**DOI:** 10.64898/2026.06.12.731840

**Authors:** Thomas Chadelaud, Agathe Brault, Martial Briand, Matthieu Barret, Armelle Darrasse

**Affiliations:** Univ Angers, Institut Agro, INRAE, IRHS, SFR QUASAV, F-49000 Angers, France

**Keywords:** Seed transmission, *Phaseolus vulgaris*, *Xanthomonas citri* pv. *fuscans*, seed microbiota, Type III Secretion System

## Abstract

Seed transmission is a critical pathway for the dispersal of phytopathogenic bacteria. This transmission can occur through three main routes: floral, internal, and external. Yet the relative contribution of individual transmission routes remains poorly characterized. Using a pathosystem based on *Xanthomonas citri* pv. *fuscans* (*Xcf*) and common bean (*Phaseolus vulgaris* cv. Flavert), we quantified the efficiency of each route. Under our experimental conditions, the vascular route was the most efficient with 25% of contaminated seeds and population sizes averaging 10^7^ CFU per contaminated seed. Deploying this experimental framework to ten seed-borne bacterial strains isolated from bean revealed that almost none transmitted to seeds through any route, or at best at low efficiency. However, most of the strains were capable of surviving and disseminating within the vascular system. A major bottleneck for seed transmission was identified for pod vascular organs colonization and the similar behavior of an *Xcf* mutant, deficient in the T3SS, suggested that plant immunity could be involved at this step. Co-inoculation of a consortium composed of the seed-borne strains with *Xcf* reduced the number of seeds contaminated by *Xcf* at the highest inoculum concentration, although other consortia members were never recovered from seeds. This suggests that the strains are recognized by the plant and trigger defense responses. These findings also raise questions about the mechanisms used by seed-associated bacteria to colonize seeds *in situ*.

## Introduction

Effective management of plant bacterial diseases relies predominantly on prophylactic approaches, given the scarcity of curative treatments available once infection is established (Sundin et al. 2016). In this context, the precise identification and characterization of primary inoculum sources constitute a prerequisite for disease management strategies (Gitaitis and Walcott 2007). Phytopathogenic bacteria can persist and proliferate across a diverse array of reservoirs within cropping systems, including weeds, arthropod vectors, infected crop residues, and contaminated seed lots, each of which may contribute differentially to pathogen dispersal and initiation of epidemy (Burdman and Walcott 2012; Chen et al. 2021; Lamichhane and Bartoli 2015; Zhao et al. 2002).

Seed is the first input in cropping systems and is therefore a widely traded commodity worldwide. Based on the ISTA Reference Pest List v15 published in January 2026, seeds of eight crop species are a pathway for 17 phytopathogenic bacteria (Denancé and Malabarba 2025). This observation is not new, since the importance of seeds as a vector for the spread of phytopathogenic agents has been documented since the late 1800s (Baker and Smith 1966). As global trade increases, so does the threat of introduction of seed-transmitted plant pathogens. Ensuring seed sanitary quality is therefore critical to avoid the emergence of plant disease in new geographic areas. As per today seed sanitary control solutions mostly consist in seed testing and destruction or disinfection of the contaminated seed lots. Seed treatments can reduce bacterial infections but come with their own set of flaws such as an important shortfall, reduced seeds viability and incomplete disinfection, especially concerning endophytic bacteria (Carisse et al. 2000; Moumni et al. 2023; Sanna et al. 2022; Temple et al. 2013; Xhemali et al. 2024).

The prevention of seed colonization is a promising avenue for the control of seedborne disease. This can be achieved in two ways: host-based resistance or biocontrol strategies. Host-based resistance requires the identification of resistance sources for breeding plants that are less susceptible to seed transmission. This approach requires a thorough understanding of the seed immune response, which can be fundamentally different from that in mature plants (Hoffmann et al. 2025; Ortega-Cuadros et al. 2022; Pagán 2022). To date, no such resistance to seed transmission of bacteria has been reported. Biocontrol strategies could exploit the interactions between microorganisms that occur during the phase of transmission to the seed. In addition to the phytopathogenic bacteria transmitted by seeds, a wide variety of commensal bacteria have been detected on seeds (Simonin et al. 2022). Inoculating the plant has already been successfully tested to introduce the endophytic bacterium *Paraburkholderia phytofirmans* PsJN into wheat, corn, soybean, and pepper seeds by floral inoculation (Mitter et al. 2017). More recently a strain of *Pantoea agglomerans* was introduced in wheat and ryegrass seeds through spike inoculation (Sanz-Puente et al. 2025). Thus, commensal seedborne strains could be used as tools for biocontrol solutions against the colonization of seeds by pathogenic bacteria (Barret et al. 2016).

Regardless of the strategy used (*i.e.* plant breeding or biocontrol-based approach), a better understanding of the transmission routes used by these microorganisms to colonize seed is needed. Seed transmission routes can be schematically divided into three pathways (Maude 1996). The first is the vascular route, where the microorganism colonizes the seed through the xylem. This route is used, for example, by *Clavibacter michiganensis* during transmission to tomato seeds (Nandi et al. 2018). For the floral route, the microorganism enters the flower and colonizes the stigma, the style until the ovary. This pathway is typically used by *Paracidovorax citrulli* during transmission to watermelon seeds (Bergmann et al. 2026; Dutta et al. 2015). Last, the external route consists in the colonization of the seed by direct contact of microorganisms with the fruit. This pathway is used by *Pseudomonas savastanoi* pv. *phaseolicola* during transmission to bean seeds (Taylor et al. 1979). These routes are not mutually exclusive. Their relative importance remains unknown; however, they each play a different role. Hence, the vascular and floral routes of contamination allow an early and internal colonization of seeds, which can ensure a better survival rate for bacteria than the external route (Darrasse et al. 2018; Dutta et al. 2012).

*Xanthomonas citri* pv. *fuscans* (*Xcf*), a causal agent for the common bacterial blight (CBB) on bean can use all three routes (Chen et al. 2021; Darsonval et al. 2008). Genetic inverse approaches highlight the role of the type III secretion system (T3SS) and adhesins in the transmission of *Xcf* to bean seeds, particularly *via* the vascular pathway (Darsonval et al. 2008, 2009). Simultaneous monitoring of the expression of *Xcf* and bean seed genes reveals a phase of active interactions between the plant and the bacteria during seed maturation, followed by a much quieter phase at maturity. This analysis results in a proposal for a “ceasefire” *scenario*, specific to seed tissues, allowing the contaminated seed to reach maturity and retain its germination capacity (Darrasse et al. 2024). Taken together, these studies suggest that plant immunity can be activated during seed formation and that bacteria capable of transmitting to seeds use their T3SS to bypass plant defenses. Since common bean has the particularity of harboring a low diversity of microorganisms as well as a small population size on a given seed (Chesneau et al. 2022), seed colonization could be competitive on this model. Thus, we used this specificity to explore the biocontrol potential of commensal strains to prevent *Xcf* colonization of bean seeds.

The first goal of this study was to monitor the relative importance of each route for the transmission of *Xcf* to common bean seeds to identify potential bottlenecks. We also extended the study of these routes to commensal bacteria from a collection of common bean seed-isolated strains (Arnault et al. 2024) and explored their potential use for biocontrol against the transmission of *Xcf* to the bean seeds.

## Material and methods

### Bacterial strains and culture media

*E. coli* DH5α was grown at 37°C in lysogeny broth (LB, 10 g liter^-1^ tryptone, 10 g liter^-1^ NaCl, 5 g liter^-1^ yeast extract) supplemented when needed with 14 g liter^-1^ of agar and 100 µg ml^-1^ of ampicillin (Amp100). Other bacterial strains (Table 1) were grown at 28°C in tryptone soy broth (hereafter TSB, 17 g liter^-1^ tryptone, 3 g liter^-1^ soybean peptone, 2.5 g liter^-1^ glucose, 5 g liter^-1^ NaCl and 5 g liter^-1^ K_2_HPO_4_) or 1/10 strength tryptone soy broth (TSB10) supplemented when needed with 14 g liter^-1^ of agar (TSA) and 50 µg ml^-1^ or 200 µg ml^-1^ of rifamycin (hereafter Rif50 or Rif200) or 50 µg ml^-1^ of kanamycin (Kan50). The growth of all strains was tested in TSB10 at 28°C using a SPECTROstar Nano (BMG LABTECH, Champigny sur Marne, France) with double orbital shaking at 300 rpm for 48 h.

**Table 1:** Bacterial strains used in the study.

|  |  | T3SS<sup>a</sup> | Origin |  |
| --- | --- | --- | --- | --- |
| Strains | Description |  | Origin tissues | /Reference |
| *Escherichia coli* |  |  |  |  |
| *E. coli* DH5α | F-, *Δ(argF-lac)169*,<br>*φ80dlacZ58(M15)*, *ΔphoA8*, *glnX44(AS)*, *λ-deoR481*, *rfbC1*, *gyrA96(NalR)*, *recA1*, *endA1*, *t*<br>*hiE1*, *hsdR17* |  |  | (Hanahan 1983) |
| *Xanthomonas citri* pv.<br>*fuscans* (*Xcf*) |  |  |  |  |
| *Xcf*CFBP 13761 | rifamycin resistant<br>variant of *Xcf*CFBP<br>7767 isolated from<br>common bean leaf | Hrp2 | *P. vulgaris* leaf | (Darrasse et al. 2024) |
| *Xcf*CFBP 13761<br>*ΔhrcV* | deletion mutant defective<br>for T3SS | Hrp2<br>*ΔhrcV* |  | this study |
| *X. campestris* pv.<br>*campestris* (*Xcc*) |  |  |  |  |
| *Xcc* CFBP 8340 | *Xcc* 8004 reference strain | Hrp2 | *Brassica oleracea* var.<br>*botrytis* leaf | (Turner et al. 1984) |
| *Kosakonia* sp. |  |  |  |  |
| *Kosakonia* sp.<br>CFBP 8986 | wild type (WT) | Hrp1 | *P. vulgaris* seed | (Arnault et al. 2024) |
| *Kosakonia* sp.<br>CFBP 8986-R | rifamycin resistant<br>variant of the WT strain | Hrp1 |  | this study |
| *Pantoea agglomerans* |  |  |  |  |
| *P. agglomerans*<br>CFBP 13583 | WT | Hrp1 | *P. vulgaris* seed | (Chesneau et al. 2020) |
| *P. agglomerans*<br>CFBP 13583-R | rifamycin resistant<br>variant of the WT strain | Hrp1 |  | this study |
| *P. agglomerans*<br>CFBP 8988 | WT | Hrp1 | *P. vulgaris* seed | (Arnault et al. 2024) |
| *P. agglomerans*<br>CFBP 8988-R | rifamycin resistant<br>variant of the WT strain | Hrp1 |  | this study |
| *P. agglomerans*<br>CFBP 9000 | WT |  | *P. vulgaris* seed | (Arnault et al. 2024) |
| *P. agglomerans*<br>CFBP 9000-R | rifamycin resistant<br>variant of the WT strain |  |  | this study |
| ***Leclercia* sp.** |  |  |  |  |
| *Leclercia* sp. CFBP<br>8987 | WT |  | *P. vulgaris* seed | (Arnault et al. 2024) |
| *Leclercia* sp. CFBP<br>8987-R | rifamycin resistant<br>variant of the WT strain |  |  | this study |
| ***Pseudomonas korrensis*** |  |  |  |  |
| *P. korrensis* CFBP<br>9003 | WT |  | *P. vulgaris* seed | (Arnault et al. 2024) |
| *P. korrensis* CFBP<br>9003-R | rifamycin resistant<br>variant of the WT strain |  |  | this study |
| ***Pseudomonas viridiflava*** |  |  |  |  |
| *P. viridiflava* CFBP<br>13578 | WT | Hrp1,<br>Rhizobi<br>ales | *P. vulgaris* seed | (Chesneau et al. 2020) |
| *P. viridiflava* CFBP<br>13578-R | rifamycin resistant<br>variant of the WT strain | Hrp1,<br>Rhizobi<br>ales |  | this study |
| *P. viridiflava* CFBP 8985 | WT | Hrp1, Rhizobiales | *P. vulgaris* seed | (Arnault et al. 2024) |
| *P. viridiflava* CFBP 8985-R | rifamycin resistant variant of the WT strain | Hrp1, Rhizobiales |  | this study |
| ***Siccibacter* sp.** |  |  |  |  |
| *Siccibacter* sp. CFBP 8990 | WT |  | *P. vulgaris* seed | (Arnault et al. 2024) |
| *Siccibacter* sp. CFBP 8990-R | rifamycin resistant variant of the WT strain |  |  | this study |
| ***Sphingomonas* sp.** |  |  |  |  |
| *Sphingomonas* sp. CFBP 9019 | WT |  | *P. vulgaris* seed | (Arnault et al. 2024) |
| *Sphingomonas* sp. CFBP 9019-R | rifamycin resistant variant of the WT strain |  |  | this study |

<sup>a</sup>Type 3 secretion system, the presence of T3SS in bacterial genome sequences was investigated with MacSyFinder v2 (Néron et al. 2023). The affiliation of these secretion systems to different T3SS families (SPI-1, SPI-2, Ysc, Chamydiales, Rhizobiales, Hrp1 and Hrp2) was inferred from a phylogeny of the orthologous group COG4791, following a procedure described earlier (Barret et al. 2013).

### Selection of rifamycin resistant variants strains

Bacteria were grown on TSA for 48 h at 28°C and colonies were resuspended in sterile distilled water at an OD_600 nm_ of 0.5. Bacterial suspensions were plated on TSA supplemented with Rif200 and grown at 28°C for 72 h to select resistant clones.

### Construction of T3SS deletion mutant *Xcf* CFBP 13761 Δ*hrcV*

The deletion mutant *Xcf* CFBP 13761 Δ*hrcV* was made using the allelic exchange « *sacB* » method (Engler et al. 2008; Logue et al. 2009; Pelicic et al. 1996). The primer pairs ΔhrcV.1F/ΔhrcV.1R and ΔhrcV.2F/ΔhrcV.2R (Suppl. S1) were used to amplify the 600 bp flanking both sides of the *hrcV* gene. PCR assays were performed in 50-μl volume containing 200 μM dNTP, 1.5 mM MgCl_2_, 0.5 μM each primer, 1 U of Taq Phusion Hot Start High-Fidelity DNA Polymerase (New England Biolabs, Evry, France), and 5 μl of a boiled bacterial suspension (approximately 1 × 10^7^ CFU ml^-1^). PCR conditions were 5 min at 94°C; followed by 35 cycles of 30 s at 94°C, 30 s at 62°C, and 40 s at 72°C; and ended with 5 min elongation at 72°C. Sizes of PCR fragments were checked by electrophoresis in 1.5% agarose gel. Fragments were purified using the kit Wizard® SV Gel and PCR Clean-Up System (Promega, Charbonnières-les-Bains, France) and dosed with a Nanodrop spectrophotometer (Thermo Fischer Scientific, Illkirch-Graffenstaden, France). Fifteen ng of each fragment were cloned in 25 ng of the pΔ13 plasmid (Suppl. Table S1) in a 10-µl volume with 5 U of *Bsa*I restriction enzyme (Fermentas, Saint-Rémi-lès-Chevreuse, France) and 2.5 U of T4 DNA ligase using the appropriate buffer (Promega). Incubation was performed at 37°C for 1 h then, 55°C for 10 min and at 80°C for 5 min. *E. coli* DH5α cells were transformed with the recombinant plasmid pΔ13*ΔhrcV* (Suppl. Table S1) using a standard heat-shock protocol (42°C for 30 sec, followed by 37°C for 1 hour in LB). The transformants were spread on LB agar supplemented with Amp100 and grown for 24 h, at 37°C. The resulting colonies were verified by PCR amplification of *hrcV* gene with the primer pair ΔhrcV.F/ΔhrcV.R (Suppl. Table S1). The purified recombinant plasmid was transformed into competent *Xcf* CFBP 13761 by electroporation at 2 kV for 5 ms in 0.2 cm cuvettes (Gene Pulser/MicroPulser Electroporation Cuvettes, 2.0 cm gap #1652086, BioRad, Marnes-la-Coquette, France) in the MicroPulser Electroporator (BioRad) followed by culture in TSB at 28°C for 2 h. The transformants were spread and selected on TSA supplemented with Kan50. The *Xcf* CFBP 13761::pΔ13Δ*hrcV* merodiploïds were counter-selected on TSA supplemented with sucrose (10% w/v) at 28°C for 48 h. The resulting colonies were spotted on TSA with or without Kan50. Kanamycin-sensible colonies were verified by PCR amplification of *hrcV* gene. *Xcf* Δ*hrcV* was verified by long read DNA sequencing (see Long read DNA extractions and sequencing section).

### Inoculum preparation

For inoculum preparation, 24 h-old bacterial cells in pure culture were collected from plates and suspended in sterile distilled water. Suspension was calibrated at OD_600 nm_ 0.1 and checked by dilution-plating to verify that it complied with the expected value of 1 × 10^8^ CFU ml^-1^ (Suppl fig 1). When necessary, inoculum was adjusted to the desired final concentration with sterile distilled water, or mixed with other inocula for co-inoculation experiments. A Bacterial consortium of 10 strains: *Kosakonia* sp. CFBP 8986-R, *Pantoea agglomerans* CFBP 13583-R, *P. agglomerans* CFBP 8988-R, *P. agglomerans* CFBP 9000-R, *Leclercia* sp. CFBP 8987-R, *Pseudomonas korrensis* CFBP 9003-R, *Pseudomonas viridiflava* CFBP 13578-R, *P. viridiflava* CFBP 8985-R, *Siccibacter* sp. CFBP 8990-R, and *Sphingomonas* sp. CFBP 9019-R was assembled with an identical volume of all bacterial suspensions at an OD_600 nm_ of 0.1. Suspensions of *Xcf* CFBP 13761 and *Xcf* CFBP 13761 *ΔhrcV* at an OD_600 nm_ of 0.1 were added at a 1:1 (v/v) ratio (*Xcf* : Consortium) to create two new bacterial mixes (*Xcf* CFBP 13761 + Consortium and *Xcf* CFBP 13761 *ΔhrcV* + Consortium). Co-inoculation experiments were done with pure or 10 to 1000 times-diluted suspensions. All inocula done in this study were plated for quality control of the experiment (**Fig. S1**).

### Culture conditions of bean plants

Seeds of common bean (*Phaseolus vulgaris*) cv. Flavert, susceptible to common bacterial blight, were provided by Vilmorin (La Ménitré, France) and stored at 9°C. Seeds were sown in one-liter pots with substrate tray (NF U 44-551, Klasmann-Deilmann GmbL, Ruy Montceau, France) in climatic chambers with 16 h of day light, 70% hygrometry, 23°C daytime/ 20°C nighttime. Plants were pinched after the third leaf. Plants were watered twice a week for the first three weeks, after which a nutrient solution with an N/P/K ratio of 15:10:30 was used. For pathogenicity experiments, the temperature was set to 28°C during the day and 25°C at night with 95% humidity the day before inoculation, when the first trifoliate leaf had fully expanded (two weeks after sowing). This temperature and humidity were maintained until the end of the assays. For seed transmission experiments, the temperature was set to 25°C during the day and 22°C at night three days before inoculation when the plants were at the flower bud stage (four weeks after sowing). This temperature was maintained until the end of the experiments.

### Pathogenicity assays

Pathogenicity of *Xcf* CFBP 13761 and *Xcf* CFBP 13761 *ΔhrcV* strains was assessed on bean (*Phaseolus vulgaris* cv. Flavert). Briefly, the first trifoliate leaf of two-week-old bean plants was immersed in bacterial suspensions at an OD_600 nm_ of 0.1 (∼1.10^8^ CFU ml^-1^) or in sterile osmosed water. After an additional two weeks of plant growth, symptoms were analyzed with Ilastik (version 1.4) and ImageJ (version 1.53).

### Endpoint seed transmission assays

Inoculations were done by infiltration with a needle of one µl of bacterial suspension at a OD_600 nm_ of 0.1 (∼10^8^ CFU ml^-1^). For vascular pathway, stem was infiltrated three cm under flower buds at pollination stage. Floral pathway was tested by either infiltrating closed flower buds three days before or on the day of pollination, or by depositing a one-µl droplet in opened flower one day after pollination (DAP). External inoculations were done by immersing pods at seven, 21, or 35 DAP in a 1/1000 dilution of bacterial suspension at an OD_600 nm_ of 0.1 (∼10^5^ CFU ml^-1^). Mock samples were inoculated in the same way using sterile osmosed water. Serial dilutions of all inocula were plated and incubated at 28°C for 72 h on TSA10 before colony counting. At 42 DAP, pods were harvested directly upstream of the inoculation marks. Seeds were extracted from pods with caution to avoid any pollution by external contact. Three seeds per inoculated pods were analyzed individually (one closed to the peduncle, one in the middle of the pod and the third one at the end of the pod) in the three experiments comparing the different route of contamination by *Xcf* CFBP 1376. In all other experiments, seeds were pooled per plant to constitute a seed lot. Each seed lot was homogenized and divided into several subsamples of three to ten seeds, except for *Xcf* CFBP 13761, for which the seeds were analyzed individually. Subsampling was performed to determine the percentage of contaminated seeds. The size of the subsample was increased to maximize the number of seeds analyzed when a low or zero contamination rate was expected. Seeds were soaked in one ml of water per seed. Seed extracts of *Xcf* CFBP 13761 were serial-diluted in sterile water and 50 µl were plated on TSA10 supplemented with Rif50. For the other strains, up to five ml of the seed extracts were directly plated on TSA10 supplemented with Rif50. Plated volumes were increased to avoid the negative impact of an increased subsample size on the detection threshold (below one CFU per plated volume). All plates were incubated at 28°C for 72 h. The percentage of contaminated seeds within the seed lot was estimated using the following formula: 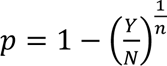. Y: number of healthy groups, N: number of analyzed groups, n: number of seeds per group (Maury et al. 1985).

### Kinetic seed transmission assays

Inoculations were done at pollination stage by infiltration of one µl of bacterial suspension at an OD_600 nm_ of 0.1 in the stem, three cm under the flower buds with a needle. Mock samples were inoculated in the same way with sterile osmosed water. At seven, 15, 21, and 35 DAP, stem and pods directly above the inoculation mark were harvested and dissected with sterile instruments. For seven and 15 DAP samples, a stem fragment located in between one and two cm upstream the inoculation mark and the whole pod were analyzed separately. Furthermore, for 21 and 35 DAP, the first seed of the pod (closest to the pod stalk), its funicle, and one-cm fragment of the ventral suture where the seed was attached were analyzed separately. Stem and seed samples were soaked whereas suture, funicle and pod samples were grounded in sterile water. Fifty µl of dilution series were plated on TSA10 supplemented with Rif50 and incubated at 28°C for 72 h.

### Vascularization assays

Stems were inoculated by infiltration one µl of a bacterial suspension at an OD_600 nm_ of 0.1 into the stem with a needle. Mock samples were inoculated in the same way with sterile osmosed water. Stem samples (from 2 cm below the inoculation mark to 4 cm upstream) were cut 72 h post-inoculation, surface sterilized with bleach (Bec Formule Pro, 2.6% hypochlorine, PLG Group, Saint Aignan de Grand Lieu, France), and divided into one-cm pieces from the point of inoculation. Stem imprints were done by pressing each stem five times onto TSA10 supplemented with Rif 50. Plates were incubated at 28°C for 72 h before numbering the colonized imprints.

### Long read DNA extractions and sequencing

DNA extraction was done using the NucleoBond HMW DNA (Macherey-Nagel, Düren, Germany) and the NucleoMag X32 system (Macherey-Nagel, Düren, Germany). Large DNA fragments were selected by purification with AMPure XP beads (Beckman Coulter, Brea, California, USA). The sequencing was done by MinION sequencing with a MinION Flow Cell (Oxford Nanopore Technologies, Oxford, UK). Sequences were assembled using Flye v2.9.5-b1801 (Kolmogorov et al. 2019). Sequence start was fixed using the fixstart option of Circlator v1.5.5 (Hunt et al. 2015). Polishing was performed using Flye v2.9.5-b1801. The assembled genomic sequence of mutant was compared with the wild-type sequence GCF_002759375.1 using SkIf2 (Denancé et al. 2019).

### Statistics

All statistics and figures were done with RStudio. The analysis of endpoint transmission percentages of contamination for *Xcf* routes, kinetic vascular transmission, vascularization, endpoint transmission percentages for seed-isolated strains for floral route were done with generalized linear model (glm) using a binomial distribution. The population size on seeds for *Xcf* transmission routes was analyzed with a glm using a Poisson distribution. Vascular and external transmission percentages for seed-isolated strains were analyzed using a Kruskal-Wallis test and associated Dunn’s post-hoc test using rstatix package (v0.7.2). Conditions of applications for the glm analysis were checked using the DHARMa package (v0.4.7) and marginal means were obtained using the emmeans package (v1.11.2-8). The figures were done using ggplot2 (4.0.0), ggbeeswarm (v0.7.2), ggbreak (v0.1.6), ggeffects (v2.3.1), ggstatsplot (v0.13.3), cowplot (v1.2.0) and ggpubr (v0.6.1) and refined with Inkscape 1.3.2.

## Results

### The vascular route is the most efficient of the transmission of *X. citri* pv. *fuscans* to bean seeds

The relative importance of the different seed transmission routes described for phytopathogenic agents (internal, floral and external) was first estimated for the bacterium *X. citri* pv. *fuscans* CFBP 13761. For the floral route, the percentage of contaminated seeds varied between 2 to 5% (**Fig. 1A**). The percentage of seeds contaminated by *Xcf* CFBP 13761 increased significantly when inoculated by the vascular pathway, with 25% of contaminated seeds, and by external pathway, with 15% to 24% of contaminated seeds depending on the inoculation date. No correlation was found between the inoculation pathway and the position of contaminated seeds in the pods. When examining the population sizes found within each contaminated seed, these ranged from 10^2^ to 10^3^ CFU per seed when *Xcf* CFBP 13761 was inoculated *via* floral and external routes (**Fig. 1B**). By contrast, vascular route allowed a significantly (padj < 0.05) higher *Xcf* CFBP 13761 population sizes on seeds with an average of 10^7^ CFU per contaminated seed. These results showed that *Xcf* CFBP 13761 transmitted to seeds through all three routes, albeit with varying degrees of efficiency. Taking into account both contamination percentages and population sizes, the vascular route was the most efficient means of seed transmission for *Xcf* CFBP 13761.

**Figure 1:**
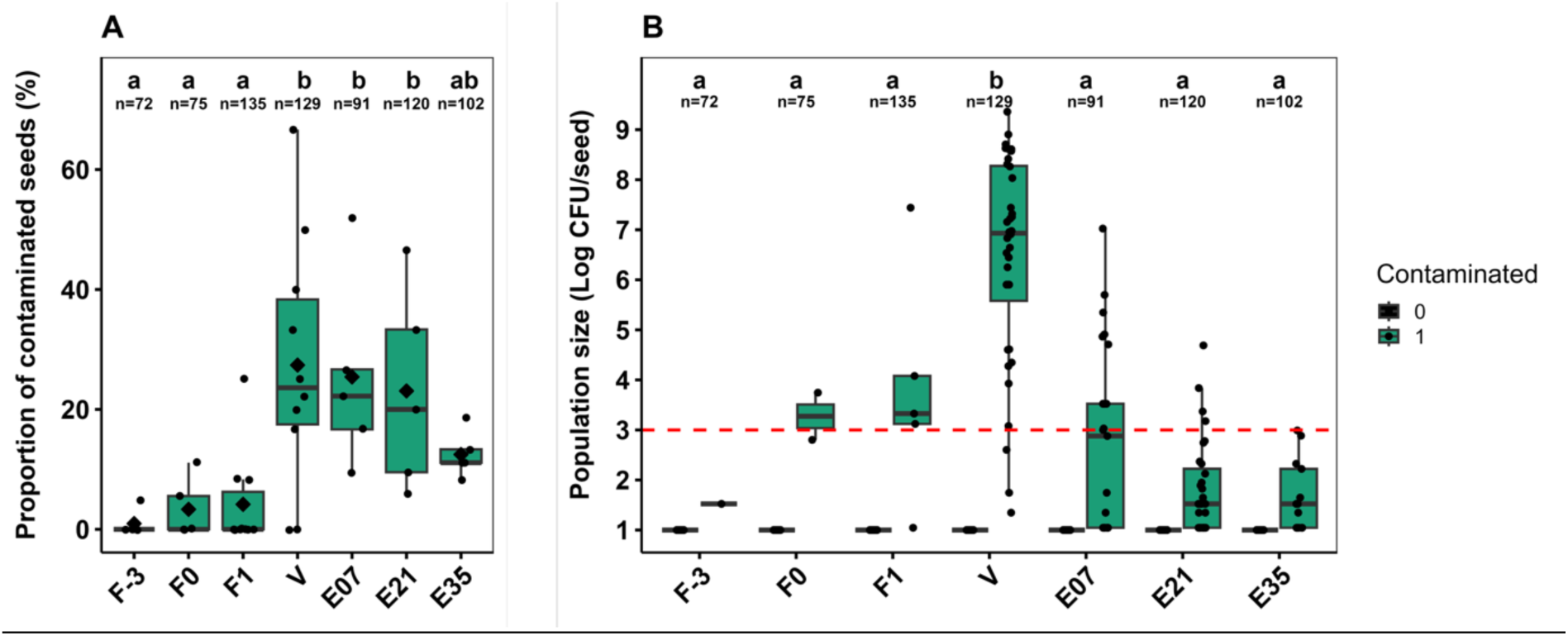
Transmission routes of *Xanthomonas citri* pv. *fuscans* to bean seeds. A. Proportion of seed contamination by *Xcf* for floral (F), vascular (V) or external (E) routes. Inoculations were performed three days before pollination (DBP, -3), at pollination (0), or one, seven, 21, 35 days after pollination (DAP). Contamination levels for each route were analyzed using a general linear model based on a binomial probability distribution. Significant groups were identified with a padj < 0.05. B. *Xcf* population size on seeds inoculated by floral (F), vascular (V) or external (E) routes. Black circles indicate the percentages of contaminated seeds in a seed lot and the diamonds represent the mean percentage for each strain. The red dotted line is the population size threshold required for an effective transmission of *Xcf* from the seed to the seedling (Darrasse et al. 2007). Population sizes for each route were analyzed using a general linear model based on a Poisson probability distribution. Significant groups were identified with a padj < 0.05. Results correspond to the cumulated results of three independent experiments for a total of 15 plants per strain with at least three separate inoculations per plant. n: number of seeds analyzed.

### The vascular transmission of bean-seed isolates to seeds is a rare event and is dependent of the T3SS

Having identified the vascular pathway as the most efficient route for *Xcf* transmission to bean seeds, we applied this method of inoculation to the nine rifamycin resistant variants of seed-isolated strains (**Table 1**). The growth of the each rifamycin resistant variant was tested to ensure that it was comparable to the corresponding WT strain (**Fig. S2**). In order to improve detection of rare events in increasing the number of analyzed seeds, we pooled all the seeds of a plant in a seed lot. Subsampling was performed to determine the number of contaminated seeds per seed lot and allowed to analyze up to 30 seeds per seed lot. To avoid a loss of detection sensitivity due to a larger soaking volume, we increased the volume of the spread sample. The strain *X. campestris* pv. *campestris* 8004, a pathogenic bacterium of the *Brassicaceae* was used as an incompatible interaction control on a non-host plant (Darsonval et al. 2008). Moreover, a T3SS-deficient mutant of *Xcf* CFBP 13761 was also inoculated as this secretion system has previously been described as a major determinant in the transmission of another strain of *Xcf* to bean seeds (Darsonval et al. 2008). The sequencing of *Xcf* CFBP 13761 *ΔhrcV* (PRJNA1477745) confirmed the deletion of the *hrcV* gene but also revealed a deletion of a portion of the *uvrB* gene, encoding the protein B of the UvrABC system, a bacterial multi-enzyme complex involved in DNA repair (Van Houten and Snowden 1993). The growth of the T3SS-deficient mutant *Xcf* CFBP 13761 *ΔhrcV* was compared to the *Xcf* CFBP 13761 WT strain (**Fig. S3A**). Importantly, this partial *uvrB* deletion does not change the deficiency of the mutant to secret type III effectors (T3Es), making it still a suitable negative control since the T3SS is already known to be implicated in *Xcf* transmission to bean seeds. Nevertheless, it is worth noting that the *uvrB* deletion may have exacerbated the negative fitness effects of the T3SS deficiency because nucleotide excision repair was also impaired. We also verified that *Xcf* CFBP 13761 *ΔhrcV* has lost its pathogenicity on bean leaves (**Fig. S3B**). Under our experimental conditions, the vascular transmission of *Xcf* CFBP 13761 WT was efficient with 14 contaminated seed lots out of 18 at an average of 50% of contaminated seeds (**Fig. 2**). In contrast the T3SS-deficient mutant (*Xcf* CFBP 13761 Δ*hrcV*) was isolated in only one subsample of ten seeds out of the 54 subsamples that were analyzed. This corresponded to one contaminated seed lot out of 18 with an estimated 4% of contaminated seeds within this lot. The strain used as control for an incompatible interaction, *Xcc* 8004, did not infect any seed lots. For most other strains isolated from bean seeds, no vascular transmission was observed. The only strains capable of contaminating some seed lots were *P. viridiflava* CFBP 13578 and *P. agglomerans* CFBP 13583 (Fig. 2), with three and one contaminated subsamples out of 18, respectively. In addition, *Xcf* CFBP 13761 and *P. viridiflava* CFBP 13578 were the only strains that induced symptoms on bean plants during the experiments (data not shown).

**Figure 2:**
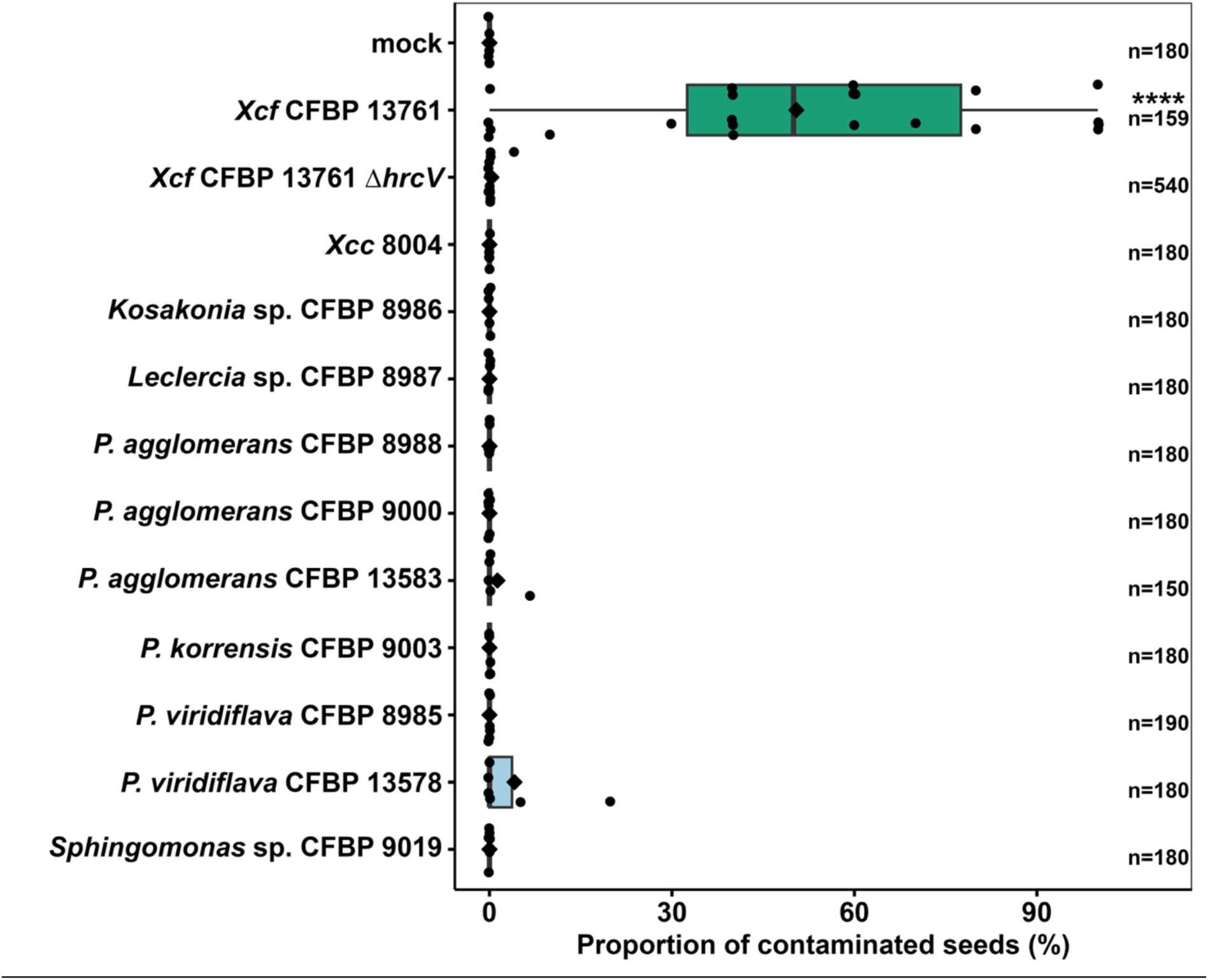
Percentage of seeds contaminated through the vascular pathway by single bacterial strains. Vascular inoculation was done at pollination stage. Black circles indicate the percentages of contaminated seeds in a seed lot and the diamonds represent the mean percentage for each strain. Contamination percentages were analyzed with Kruskal-Wallis and associated Dunn post-hoc tests. Percentages of contamination were considered significatively different for padj < 0.05. Results correspond to the cumulated results of three independent experiments for a total of 6 to 18 plants per strain with at least 3 separate inoculations per plant. n: number of seeds analyzed.

### Most seed-isolated bacteria can survive and disseminate in the bean stem

The near absence of vascular transmission of the seed-borne bacterial strains could be a result of the inoculation method within the stem. The ability of the strains to survive in the stem 72 h after inoculation was therefore evaluated. Three colonization profiles were observed (**Fig. 3**). The first profile included *Xcf* CFBP 13761, *Xcf* CFPB 13761 *ΔhrcV*, *P. agglomerans* CFPB 8988-R, *P. viridiflava* CFPB 8985-R, and *P. viridiflava* CFPB 13578-R. These strains were re-isolated in 65% to 95% of stem imprints with no significant differences between the different distances of sampling. Therefore, these strains were able of surviving but also of spreading effectively through the stem. The second profile included *Kosakonia* sp. CFPB 8986-R, *P. agglomerans* CFPB 9000-R, *P. agglomerans* CFPB 13583-R, *P. koreensis* CFPB 9003-R, and *Siccibacter* sp. CFPB 8990-R. These strains contaminated proximal sample points at the same frequency as strains of profile 1 (average percentage of 60% to 95%). However, they spread less effectively in the stem, with 20% to 55% of contaminated imprints 3 cm above inoculation mark (**Fig. 3**). The last profile included *Leclercia* sp. CFPB 8987-R and *Sphingomonas* sp. CFPB 9019-R. These two strains had a lower survival rate in the stem. This is especially evident for *Sphingomonas* sp. CFPB 9019-R with less than 10% of contaminated imprints at each distance tested whereas *Leclercia* sp. CFPB 8987-R could survive the initial inoculation with 30% to 40% of contamination of proximal sample points but failed to disseminate through the stem with less than 10% of contamination of the most distant sample points (**Fig. 3**). In brief, most of the tested strains survived and dispersed effectively in the stem, which indicate that stem colonization was not an important bottleneck for vascular transmission to bean seeds.

**Figure 3:**
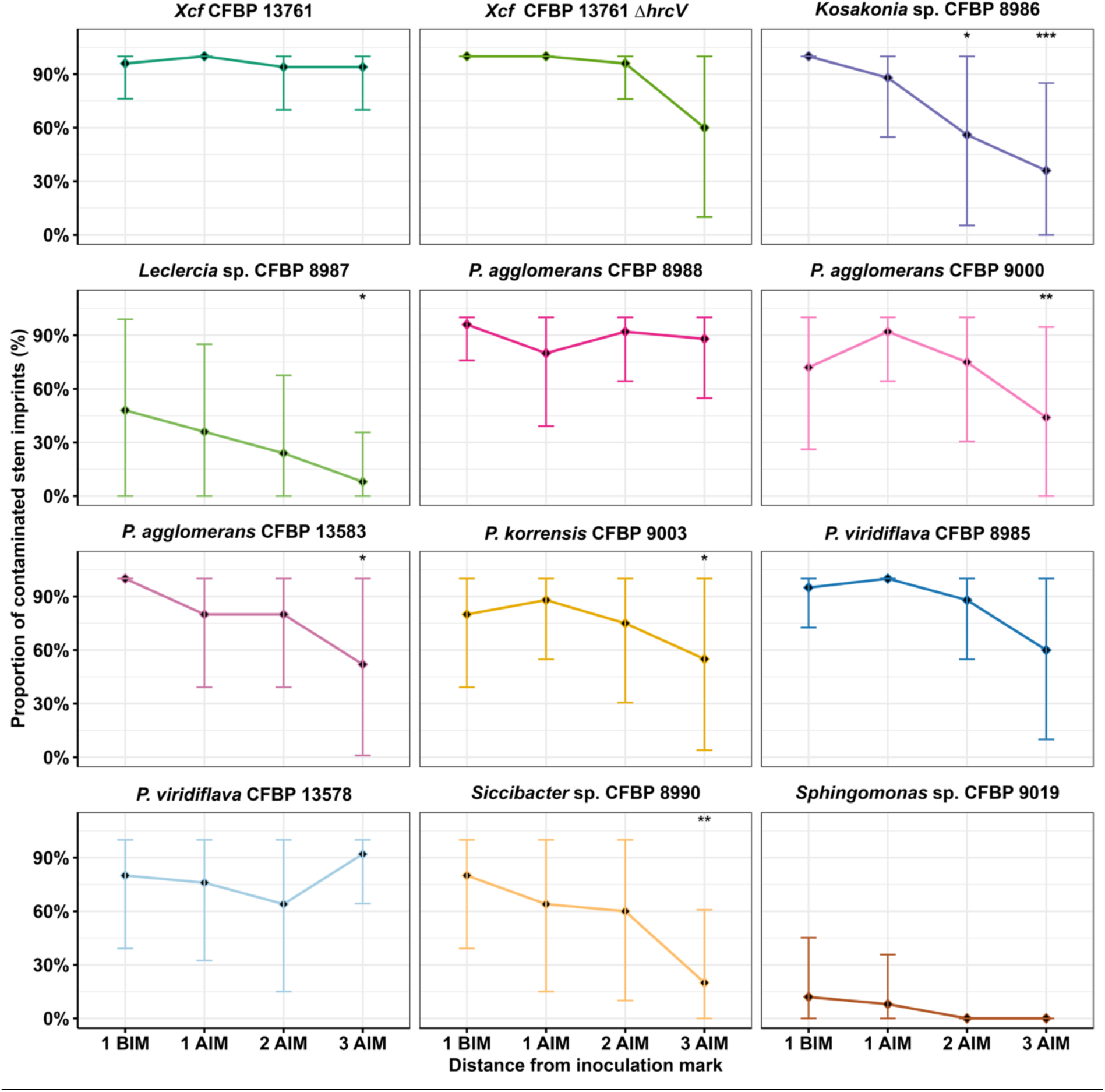
Contamination and dissemination of seed-isolated bacteria in common bean stems. Each distance corresponds to the distance of sampling from the inoculation mark; BIM and AIM corresponding to below and above inoculation mark, respectively. Contamination rates at each sampling distance were compared for each strain using general linear model based on a binomial probability distribution. *: padj < 0.1, **: padj < 0.05, ***: padj < 0.01. The experiment was performed on five plants per strain with five imprints per analyzed stem fragment.

### Pod vascular tissues colonization is an important bottleneck for the transmission to seeds

As most of the tested strains were found to survive in the stem of common bean, we explored the colonization and dissemination of the strains throughout different organs from the stem to the seed. We analyzed the stem and the young whole pod at seven and 15 DAP. Then at 21 and 35 DAP, we analyzed the stem and extracted for analysis the first seed, its funicle and one-cm long fragment of the ventral suture. To monitor the colonization of the stem, early pods, funicles, sutures and seeds, we selected a subset of strains that can survive and progress in the stem. Consistently with the previous experiment every strain had a good contamination rate of the stem above the inoculation mark, comprised between 100% and 60% at seven DAP (**Fig. 4**). This survival rate remained stable for 35 days (between 40% and 75% of contaminated samples) with no significant difference between strains. *Xcf* CFBP 13761 colonized the pods at seven and 15 DAP with 30% and 60% of samples contaminated, respectively. At later stages, *Xcf* CFBP 13761 was recovered in 70% of the sutures; 100% (21 DAP) and 70% (35 DAP) of funicles and 80% (21 DAP) and 75% (35 DAP) of seeds. Therefore, *Xcf* CFBP 13761 was capable of colonizing all the organs containing vascular vessels. In contrast, *Xcf* CFBP 13761 *ΔhrcV* only contaminated 10% of the early pods at seven and 15 DAP and failed to contaminate sutures, funicles or seeds at the latter stages. *Kosakonia* CFBP 8986 and *P. agglomerans* CFBP 8988 were unable to colonize vascular tissues beyond the stem (**Fig. 4**). *P. viridiflava* CFBP 8985 was found in 10% (seven DAP) and 20% of pods (15 DAP), but not in the different pod tissues at 21 DAP. However, it did contaminate 30% of sutures and funicles at 35 DAP without being isolated from the seeds.

**Figure 4:**
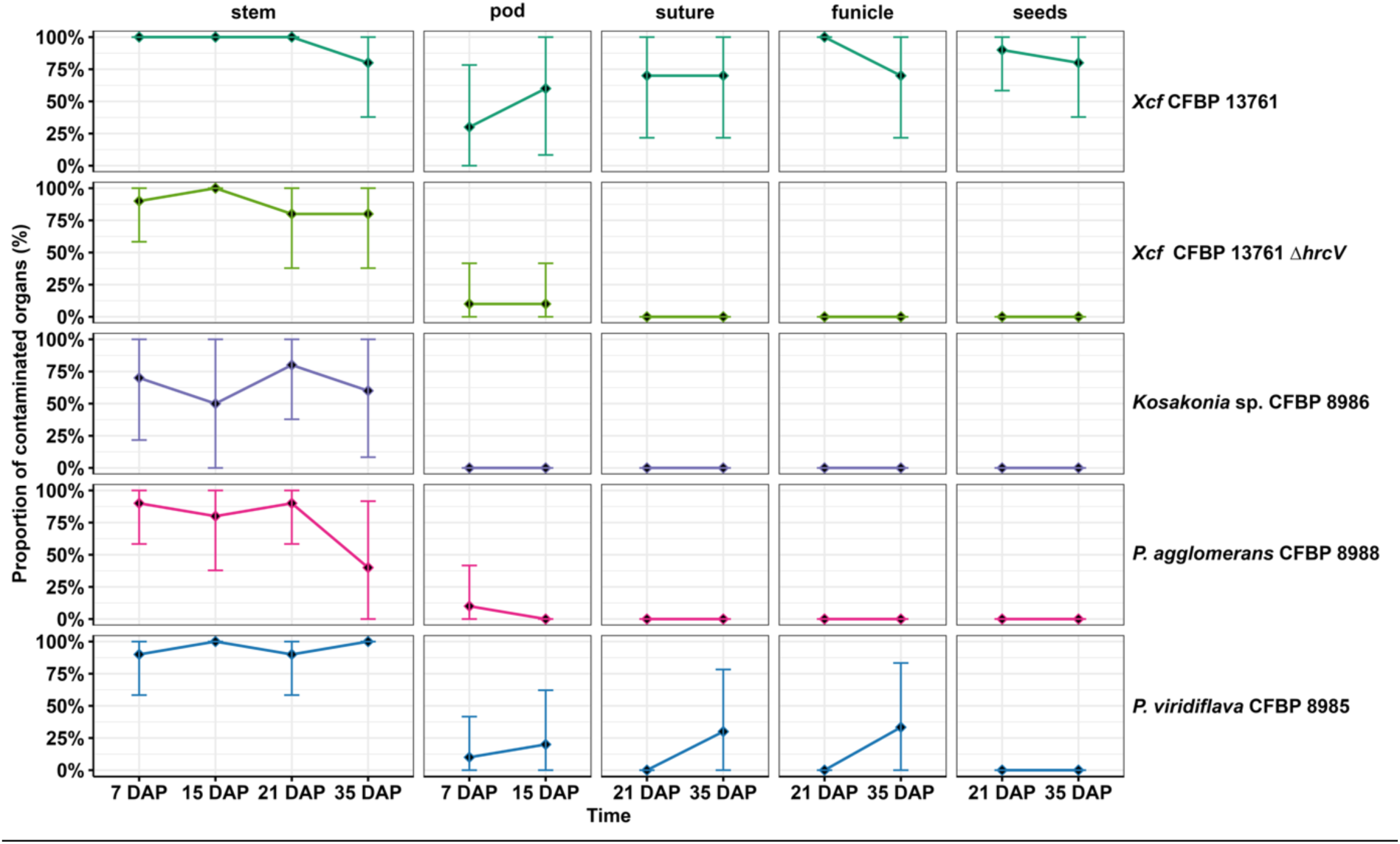
Kinetic of vascular transmission of single seed-isolated bacteria from stem to seeds. Each date corresponds to the number of days after pollination (DAP) at sampling time. Inoculation was done at pollination stage. Contamination rates at each stage were compared for each strain using general linear model based on a binomial probability distribution. Significance groups were made with a padj < 0.05. The experiment was performed on five plants with at least three separate inoculations per plant for each strain.

### The floral and external route are not sufficient for the transmission routes of bean-seed isolates to seeds

The lack of vascular transmission of most strains isolated from bean seeds under our experimental conditions may be due to the transmission route used. We next explored seed transmission by external and floral route. As previously observed, *Xcf* CFBP 13761 was able to colonize the seeds by the external route when inoculated at seven DAP with a mean 25% of seeds contaminated (**Fig. 5**). Later inoculation at 21 and 35 DAP, decreased the transmission rate to 10% and 5% of contaminated seeds. We also confirmed that floral route was the less efficient with only 5% of seeds contaminated by *Xcf* CFBP 13761 (**Fig. 5**). Almost all of the other strains tested, including the T3SS-deficient mutant of *Xcf* CFBP 13761, were unable to transmit to seeds *via* the floral and external routes (**Fig. 5**). The only exception was *Kosakonia* CFBP 8986 which contaminated two out of ten seed lots by external route when inoculated at seven DAP with contamination rates at 4% and higher than 10%.

**Figure 5:**
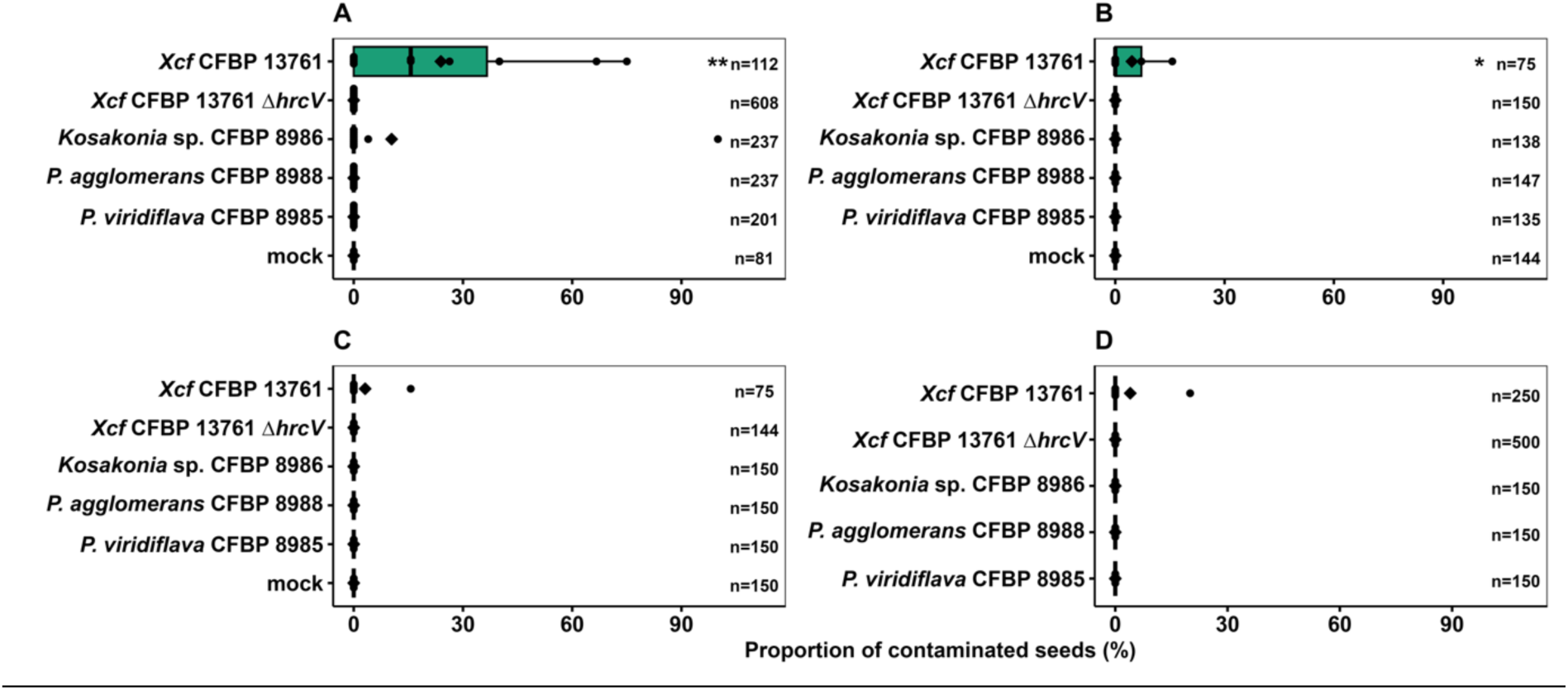
Transmission of single seed-isolated bacteria to seeds by external and floral routes. A. Proportion of seeds contaminated by single bacterial strains after inoculation by external pathway seven days after pollination (DAP). B. Proportion of seeds contaminated by single bacterial strains after inoculation by external pathway 21 DAP. C. Proportion of seeds contaminated by single bacterial strains after inoculation by external pathway 35 DAP. D. Proportion of seeds contaminated by single bacterial strains after inoculation by floral pathway at pollination. Mature seeds were sampled at 42 DAP. Black circles indicate the percentages of contaminated seeds in a seed lot and the diamonds represent the mean percentage for each strain. Contaminations rates for external route conditions were compared using non-parametric Kruskal-Wallis test and associated Dunn post-hoc test. Padj method Benjamini-Hochberg was used. Contaminations rates for floral route conditions were compared using a general linear model based on a binomial probability distribution. *: padj < 0.1, **: padj < 0.05; n: number of seeds analysezed. Cumulated results of three experiments, each performed with five plants per inoculation method and strain (with at least three separate inoculations per plant) are presented here: floral pathway was tested on 5 plants; external pathway on 10 plants for 21 and 35 DAP and on 15 plants for seven DAP.n: number of seeds analyzed.

### The use of a bacterial consortium does not improve the transmission of bean-seed isolates but impairs *Xcf* transmission rates

Failure of seed transmission for single seed-isolated bacteria may reflect recognition by the host plant, leading to the induction of plant immune responses that were not suppressed in the absence of specific type III effectors (T3Es). To investigate these hypotheses, all seed-isolated strains were assembled into a bacterial consortium and delivered *via* the vascular route. In a complementary experiment the consortium was co-inoculated with *Xcf* CFBP13761 to evaluate whether this strain could exert functional complementation through its T3Es, thereby suppressing plant defenses and enabling transmission of the consortium members. As a control a consortium was also co-inoculated the T3SS deletion mutant *Xcf* CFBP 13761 *ΔhrcV*. To test whether inoculum density influenced transmission outcomes, all inocula were serial diluted ranging from undiluted suspension to 10^-3^ dilutions. The different suspensions were verified by dilution plating and colony morphology after 72h of growth (**Fig. S1**).

As previously observed, *Xcf* CFBP 13761 was able to colonize the seeds with a mean 40% of contaminated seeds (**Fig 6**). Unfortunately, we did not detect any contamination in the seed lots inoculated with the consortium, no matter the concentration of the inoculum. The same observation applies to the co-inoculation of the consortium and *Xcf* CFBP 13761 as we only isolated *Xcf* CFBP 13761 on the plates (according to morphological observation). The transmission efficiency of *Xcf* CFBP 13761 was significantly altered when co-inoculated with the consortium at 10^5^ CFU (undiluted condition). However, this inhibitory effect was alleviated at 10^-1^ and 10^-2^ dilutions, where *Xcf* CFBP 13761 transmission rates recovered to levels comparable with lone *Xcf* CFBP 13761 inoculation. At the highest dilution (10^-3^) co-inoculation resulted in reduced transmission efficiency with only one contaminated seed detected out of 30 analyzed. These variations of *Xcf* CFBP 13761 transmission to seeds efficiency did not impact the population sizes found on contaminated seeds (**Fig. S4**). As expected, the co-inoculation on the consortium and *Xcf* CFBP 13761 *ΔhrcV* did not allow any transmission.

**Figure 6:**
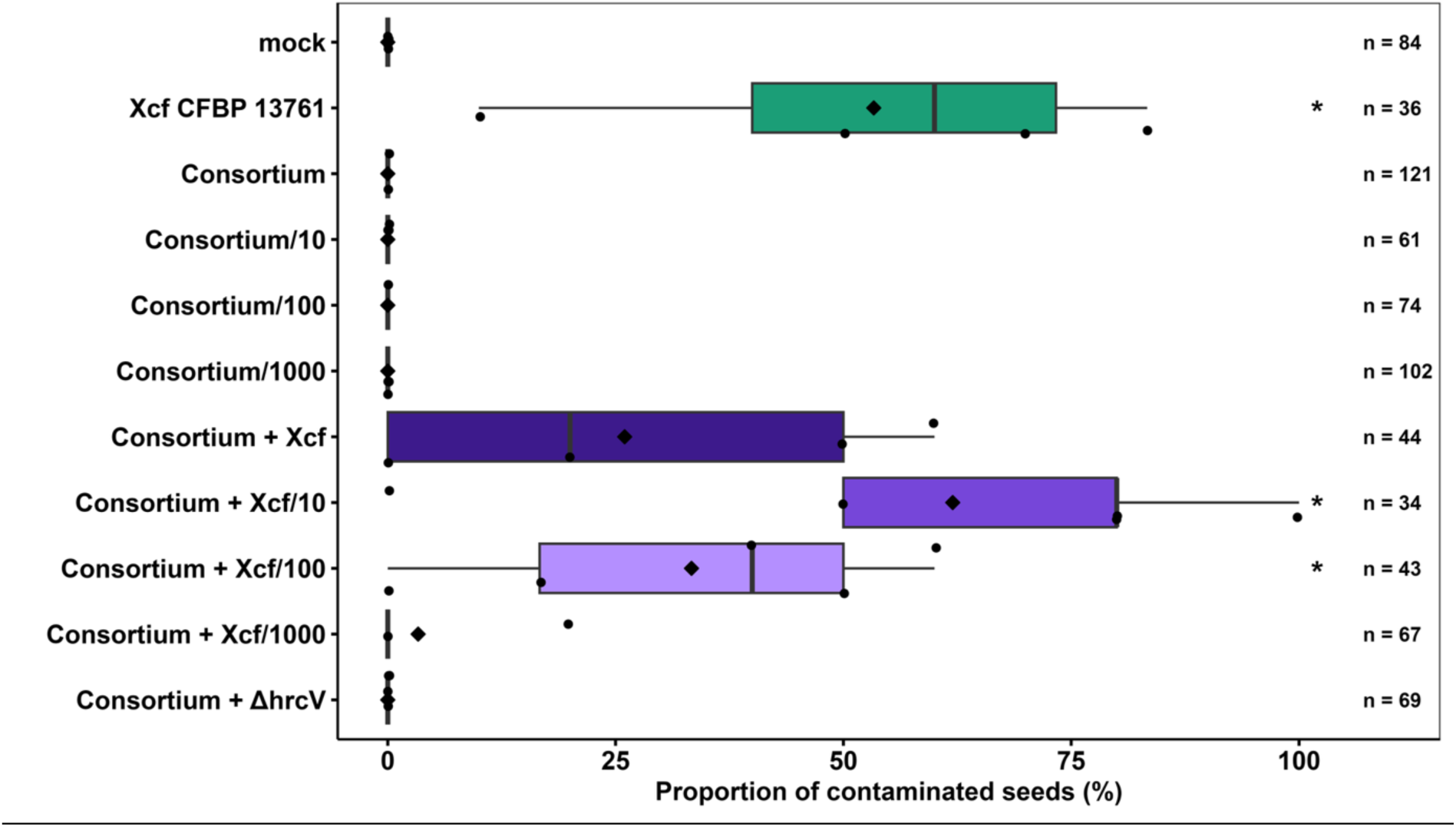
Transmission of *Xcf* co-inoculated with bean seeds-isolated bacteria to seeds by the vascular route. The Consortium condition was done by the combination of 10 bacterial strains isolated from bean seeds (see material and methods section). Consortium/10, Consortium/100 and Consortium/1000 inoculations were done with serial 1:10 dilutions of the initial Consortium. The Consortium + *Xcf* condition was done with a 1:1 mix of *Xcf* CFBP 13761 and the initial Consortium and was further serial diluted at a 1:10 ratio to do the Consortium + *Xcf* /10, Consortium + *Xcf* /100, Consortium + *Xcf* /1000 conditions. The Consortium + ΔhrcV condition was done with a 1:1 mix of *Xcf* CFBP 13761 *ΔhrcV* and the initial Consortium. *Xcf* CFBP 13761 condition was done by the inoculation of a suspension of *Xcf* CFBP 13761. Mock condition was done with the inoculation of sterile osmosed water. Inoculation was performed at pollination stage. Mature seeds were sampled at 42 DAP. Contaminations rates were compared using non-parametric Kruskal-Wallis test and associated Dunn post-hoc test. Padj method Benjamini-Hochberg was used. *: padj < 0.05; n: number of seeds analyzed. The experiment was performed on five plants per treatment with at least three separate inoculations per plant.

## Discussion

In the study, a functional pathosystem for *Xanthomonas citri* pv. *fuscans* (*Xcf*) and *Phaseolus vulgaris* was established to investigate the relative importance of the floral, vascular and external transmission routes of *Xcf* to common bean seeds. We demonstrated that the floral route is the less efficient allowing only a small fraction of seeds to be contaminated by *Xcf*. The vascular and external routes displayed similar percentages of contaminated seeds. However, the vascular route resulted in significant larger *Xcf* population sizes on seeds, making it the most efficient route for transmission. When we transposed this experimental system to other bacteria isolated from common bean seeds, none of them were consistently recovered from seed macerates, regardless of the transmission route used. The co-inoculation of *Xcf* with a consortium of seed-isolated bacteria reduced *Xcf* transmission efficiency even if consortium strains were not retrieved from the seeds. Vascular colonization assays revealed that most strains survived and disseminated within bean stem for up to 35 days post-inoculation, but failed to colonize the early pods, sutures, funicles, or seeds, with the exception of *Xcf*. Colonization of pod tissues could be an important bottleneck for seed transmission, likely involving the plant immune response since a T3SS-defective mutant of *Xcf* was unable to colonize pods or seeds.

The lower efficiency of *Xcf* CFBP 13761 to colonize bean seeds through the floral pathway compared to the external and vascular route may reflect a lesser adaptation of bean to flower-mediated infection. In watermelon, *Paracidovorax citrulli* transmitted to seeds via the floral route thanks to pollen tubes, which facilitate bacterial translocation to the ovules (Dutta et al. 2015). As common bean can undergo self-pollination while the flower is still closed (Webster et al. 1977), the stigma may not be exposed to *Xcf* CFBP 13761 at the critical developmental window, and is thus less adapted to the pathogen transmission compared to the external and vascular routes. Furthermore, in our experiments, inoculation by injection of one µl of bacterial suspension within flower buds at pollination may not sufficiently address the stigma compared to experiments performed on open flowers, such as cucurbits (Bergmann et al. 2026). However, as pollinating insects are known to improve both yield and allogamy in bean (Chacón-Sánchez et al. 2021), we cannot exclude that in field conditions, insect-pollinated flowers can be a better pathway for microbes to reach bean seeds.

Vascular and external pathways produced comparable contamination rates, however, the vascular pathway supported significantly larger population sizes on seeds. This difference may be attributed to the timing of bacterial arrival on seeds. Bacteria colonizing via the external pathway may arrive at later seed developmental stage than via the vascular pathway, when nutritional resources available for bacterial growth are less important. For instance, fluctuation of nutrient levels in clover seeds during maturation influences microbiome composition (Ahmed et al. 2025). Furthermore, decrease of water content during seed maturation could also limit bacterial multiplication (Gerna et al. 2025). Alternatively, the lowest *Xcf* population size observed on seeds *via* the external pathway may be related to a spatial pattern of colonization with bacterial cells restricted to the seed surface or in peripheral tissues (*e.g.* testa). Infections of watermelon seeds by *P. citrulli* through the pericarp are mainly located in the peripheral tissues (testa, perisperm, and endosperm) whereas they are located in the embryos for the floral pathway, but with no differences in the population sizes for both locations but with lower survival of bacteria in seeds than for floral pathway (Dutta et al. 2012).

The transmission efficiency of bean seed isolated bacteria was markedly lower with only a few seed lots contaminated across all routes tested. These results are surprising given that these bacteria were isolated from common bean seeds and are established members of the bean seed microbiota (Arnault et al. 2024; Chesneau et al. 2022; Simonin et al. 2022); although these strains, inoculated in consortia on bean seeds, colonize seedlings but are not recovered in different plant compartments at later developmental stages including seeds (Arnault et al. 2025). In our conditions, while elevated inoculum concentration may trigger plant defenses and impair seed transmission, the inoculation of diluted consortium did not allow higher transmission efficiency. The lower isolation frequency in our study relative to field-based observation (Chesneau et al. 2022) can be attributed to differences in bean culture conditions (climatic chambers versus field). The field allowed continuous plant exposure to the surrounding bacterial community, substantially increasing the probability of successful seed transmission. Moreover, our experimental system was tailored for estimating the relative contribution of each pathway in *Xcf* seed transmission. Since the three transmission routes tested (floral, internal, external) are not mutually exclusives, it is possible that the concurrent use of these three routes could result in an increase in the transmission rate. For example, spray inoculation of whole bean plants at flower bud stage, that includes all transmission routes (except contact during seed extraction), allows the transmission of *Xcc* 8004 in 20% of seed lots (Darsonval et al. 2008) and with 0.6% to 1.4% of contaminated seeds (Darrasse et al. 2010). Inoculation by combined routes (spray or immersion) also allowed the transmission of bacteria to the seeds for maize, pepper, soy, wheat and ryegrass (Mitter et al. 2017; Sanz-Puente et al. 2025). Furthermore, the reproductive type of the plant could be of great importance for the contribution of the floral route in the transmission to seeds. For example, in watermelon -a strictly allogamous species-, both pathogenic and commensal bacteria can transmit to seed by the floral route (Bergmann et al. 2026; Dutta et al. 2015). Additionally, spray inoculation at flowering stage of *P. agglomerans* did not result in seed transmission on *Arabidopsis thaliana* - self-pollinating in closed flowers like common bean- but did on ryegrass - wind-pollinated- (Sanz-Puente et al. 2025). Concerning the vascular pathway, most of the bean seed-isolated strains survived and disseminated within the stem at least three days post-inoculation. Colonization of pod vascular tissues appeared to represent a major bottleneck for seed transmission. *Pseudomonas viridiflava* CFBP 8985 was recovered from sutures and funicles at 35 DAP but was not isolated from seeds, suggesting that the transition from the funicle to the seed itself may constitute a second bottleneck.

While *Xcf* CFBP 13761 contaminated seeds via all three routes, the T3SS-defective mutant *Xcf* CFBP 13761 *ΔhrcV* was greatly impaired in its transmission ability. These results are consistent with the initial characterization of T3SS-dependent seed transmission of *Xcf* (Darsonval et al. 2008) and with subsequent studies demonstrating a role for the T3SS in suppressing plant immune responses during transmission (Darrasse et al. 2024; Terrasson et al. 2015). The inability of *Xcc* 8004 to colonize bean seeds through the vascular route, despite possessing a functional T3SS, suggests that this is because *Xcc* lacks the right T3Es. Indeed, *Xcc* T3E repertoire strongly differs from that of *Xcf* (Hajri et al. 2009). In compatible interactions, such as *Xanthomonas alfalfae* subsp. *alfalfae* with *Medicago truncatula*, defense genes - induced in seeds by *Xcc* in an incompatible interaction with *M. truncatula* - are not induced (Terrasson et al. 2015). The absence of seed contamination by *Xcc* 8004 on common bean is therefore consistent with an incompatible interaction driven by differences in the T3E repertoire.

Interestingly, the vascular monitoring of *Xcf* CFBP 13761 and the T3SS defective mutant *Xcf* CFBP 13761 *ΔhrcV* showed that the immune response does not play an important role in the stem vascular tissues colonization. However, the T3SS mutant could not colonize efficiently the early pods, sutures or funicles. Similar results have been reported for *Ralstonia solanacearum* mutants impaired in the T3SS by an insertion in *hrcV*. T3SS mutants are able to colonize xylem vessels of tomato roots and stem but not fruit peduncles (Frey et al. 1994). This result highlights the importance of an active T3SS, and by extent the role of the plant immune system, in vascular pod colonization compared to stem colonization. It should be noted that the impaired pod colonization observed in our *Xcf* mutant may be attributable to the combined effects of both the T3SS deficiency (*hrcV* deletion) and the nucleotide excision repair deficiency (*uvrB* deletion). The *uvrB* gene encodes a key component of the nucleotide excision repair (NER) pathway, which is critical for removing DNA lesions. Loss of *uvrB* function typically results in impaired DNA repair capacity and ultimately increased susceptibility to DNA-damaging agents (Seeley and Grossman 1989). When the *Xcf ΔhrcV* mutant colonizes plant tissues, it likely encounters multiple sources of genotoxic stress, including reactive oxygen species (ROS) generated by the plant immune response. These conditions would place heightened demands on the DNA reparation systems. It has been shown that the deficiency of *uvrB* lowers the mutation frequency under oxidative stress exposure (Hori et al. 2007). Yet, the role of *uvrB* in the interactions with the host remains unknown. Consequently, the combined loss of both T3SS function (preventing effective suppression of plant defenses) and NER capacity (limiting DNA repair functions) could severely compromise the mutant’s ability to survive and proliferate within plant tissues, particularly in the more immunologically active pod tissues where oxidative stress may be particularly pronounced.

As immune response control appeared to be an important factor for the transmission to the seed, we tested the hypothesis that *Xcf* could suppress the plant defenses for the bacterial community. It has been shown that *P. syringae* involves cell-to-cell differences in expression of its T3SS within plant tissue and preferentially activates its T3SS when in direct contact with apoplast cells (López-Pagán et al. 2025). Bacterial consortium transmission did not show any improvement in this matter but the presence of the consortium did hinder the transmission efficiency of *Xcf* to the seeds. We hypothesize that the presence of high quantity of commensal bacteria did prime the plant defenses to a point where *Xcf* could not contain them as well as previously observed (Darrasse et al. 2024). This could correspond to defense priming of the plant by transient microorganisms (Suteau et al. 2025). For the lower inoculum concentrations, we believe that *Xcf* can maintain the control of the plant defenses as the consortium is less concentrated, thus less activates the plant defenses. For biocontrol solutions, such transient effects are interesting as biocontrol agents that do not colonize the plant are less of a threat for biosafety. However, the absence of colonization of the seed might only allow for small effect such as seen in this study. More important effects could be obtained by inoculating seeds prior to the sowing. This method of inoculation allows for a more efficient transmission (Arnault et al. 2024) and was already used with *Stenotrophomonas rhizophila* CFBP 13503 inoculated on radish seed to reduce the colonization of the seedling by *Xcc* 8004 (Garin et al. 2024).

Our future prospects are to characterize bottlenecks of seed colonization in order to understand how many bacterial cells can enter a seed and if bacteria can multiply once within the seed and to identify which bacterial molecular determinants other than T3Es might facilitate the transmission of non-pathogenic bacteria to seeds.

## Acknowledgement

Zian Acker, Céline Moulevrier, and Anaïs Hardouin, for technical assistance. PHENOTIC Angers for Plant Phenotyping Facility, INRAE (PHENOTIC 2023), ANAN platform, SFR Quasav for DNA sequencing, CIRM-CFBP, French Collection for Plant-associated Bacteria (Perrine Portier 2017), T.C., A.B., M.B., M.B., and A.D. were supported by the third Program for Future Investments (France 2030) through the SUCSEED project (ANR-20-PCPA-0009) funded by the “Growing and Protecting Crops Differently” French Priority Research Program (PPR-CPA) under the national investment plan coordinated by the French National Research Agency (ANR).

