## Supplemental data for "The efficiency of different transmission routes of *Xanthomonas citri* pv. *fuscans* and other seed-borne bacteria to bean seeds"

**Table S1: Plasmids and primers used in this study**

**Supplementary Table S1. Plasmids and primers used in this study**

| **Plasmids** | **Description** | **Purpose** | **Reference** |
|---|---|---|---|
| pΔ13 | GoldenGate cloning compatible sacB-containing mobilizable suicide vector; Ampr | deletion mutagenesis | (Guy et al. 2013) |
| pΔ13Δ*hrcV* | recombinant pΔ13 plasmid containing the 600 pb flanking *hrcV* gene of *X.citri* pv. *fuscans* CFBP 13761 | deletion mutagenesis | this study |
| **Primers** | **Sequence** | **Purpose** | **Description** |
| ΔhrcV.F | TCATCGGGCTGATGATCCTG | deletion | *hrcV* gene |
| ΔhrcV.R | GCGGCGTGCCTCATCTGCGG | confirmation | amplification |
| ΔhrcV.1F | tttggtctcaaggtCCCAGATTGCCTGGCGGACG | cloning | 600 bp before *hrcV* |
| ΔhrcV.1R | tttggtctcacataGGCAGAGCTCCATCGCCGTC | | gene amplification |
| ΔhrcV.2F | tttggtctcatatgTGCGCAAGCCGCCCCTCCG | cloning | 600 bp after *hrcV* |
| ΔhrcV.2R | tttggtctcacttgCCTCCAGACGCTGCCTTCCG | | gene amplification |

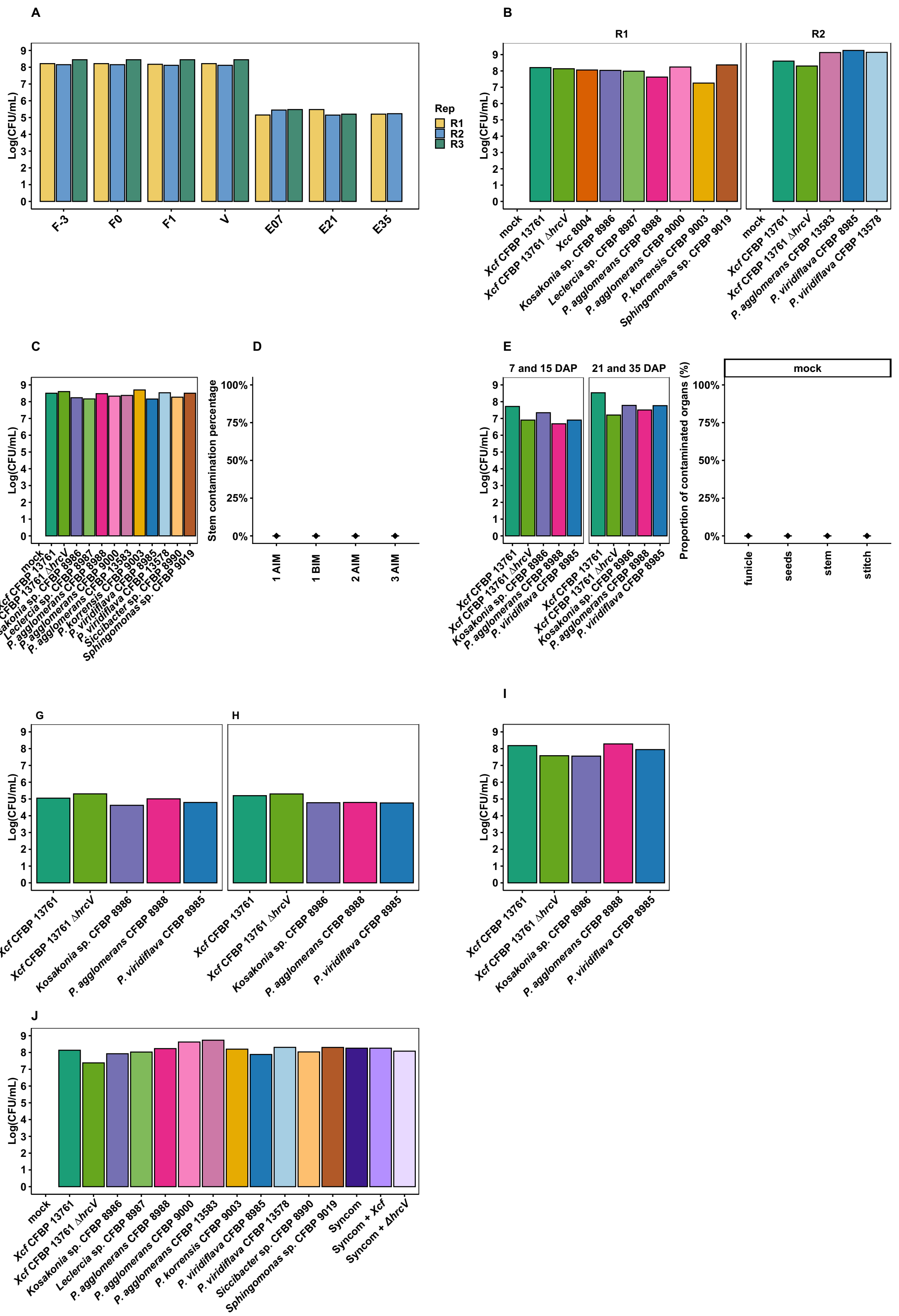


### Fig S1: Inoculum and mock controls for the transmission assays performed in the study.

A. Inoculum control for the transmission routes of *Xcf* to seeds assays. B. Inoculum control for the endpoint vascular transmission to seeds of lone inoculated bacteria. C. Inoculum control for the vascular survival of seeds-isolated bacteria in the common bean stem. D. Percentage of contamination of stem imprints 72 hours inoculated with sterile osmosed water for the vascular survival of seeds-isolated bacteria in the common bean stem assay. Each distance corresponds to the distance of sampling above the inoculation point (AIM) or below (BIM). E. Inoculum control for the kinetic monitoring of the vascular transmission of seed-isolated bacteria from common bean stem to the seed. F. Percentage of contamination of organs inoculated with sterile osmosed water during the vascular transmission of seed-isolated bacteria from common bean stem to the seed assay. G. Inoculum control for the external transmission of seed-isolated bacteria to seeds inoculated at seven and 15 days after pollination (DAP). H. Inoculum control for the external transmission of seed-isolated bacteria to seeds inoculated at 21 and 35 days after pollination (DAP). I. Inoculum control for the floral transmission of seed isolated bacteria to seeds inoculated at one DAP. J. Consortium composition and inocula control for the vascular transmission of co-inoculated seed-isolated bacteria to seeds assay. All inocula were carefully checked to contain the expected bacterial morphology in expected proportions. All inocula were plated and grown at 28°C for 72h.

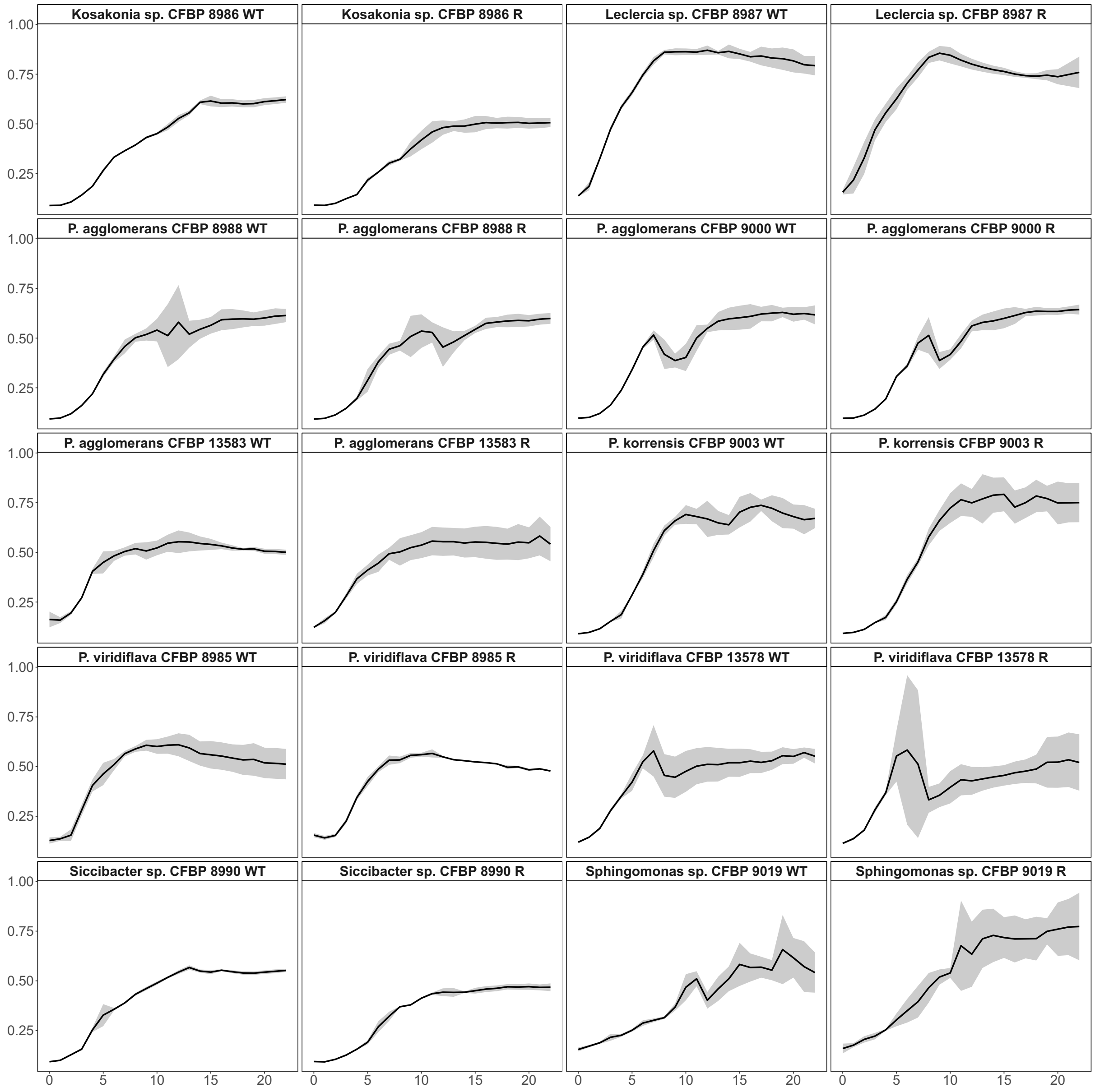


**Figure S2: Growth of the bean seeds isolated bacteria and the corresponding rifamycin resistant variants used in this study measured at $OD_{600\ nm}$.**

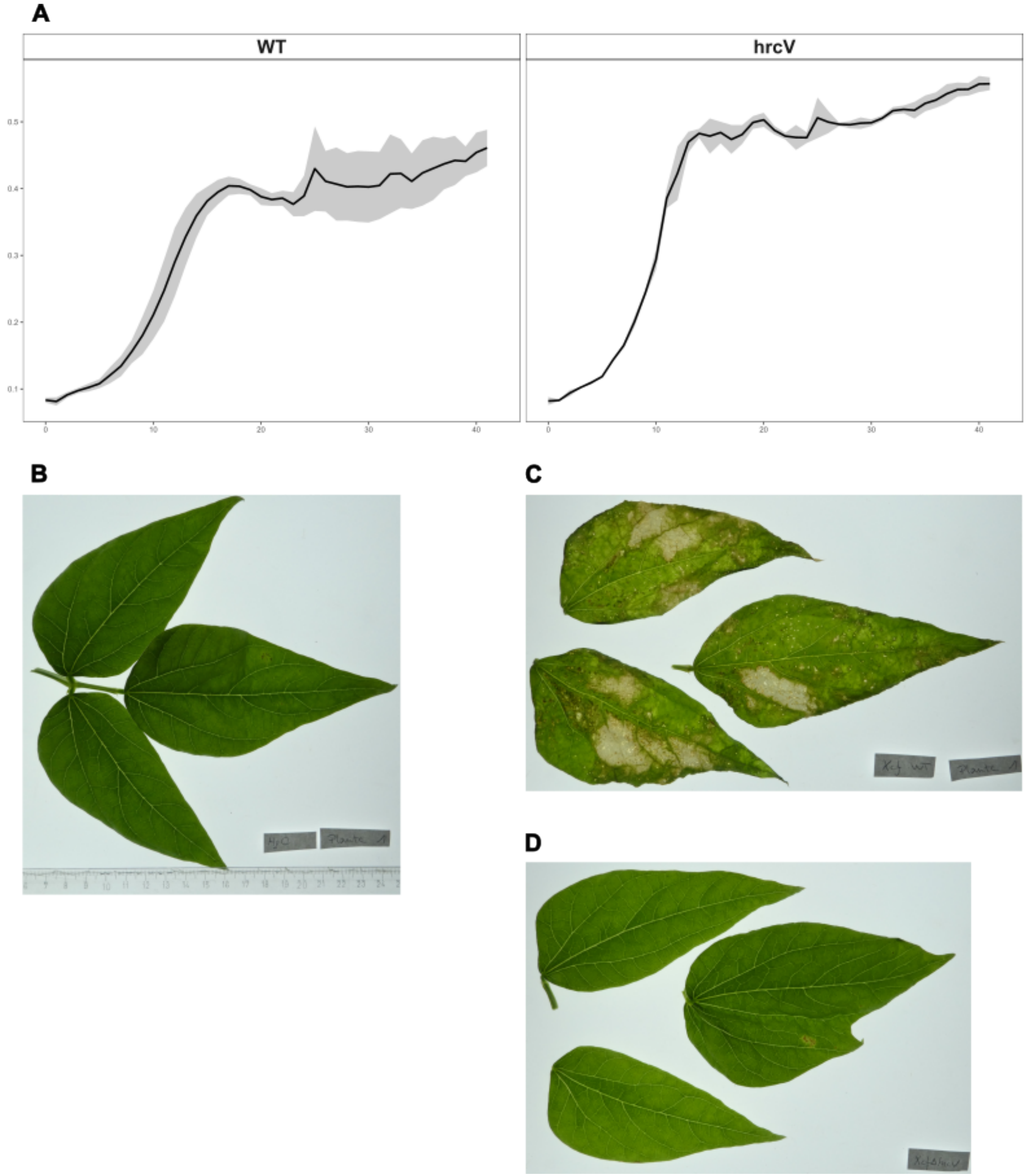


**Figure S3: A. Growth of *Xcf* CFBP 13761 WT and the deletion mutant *Xcf* CFBP 13761 *ΔhrcV* used in this study measured by $OD_{600\ nm}$. B, C and D. Pathogenicity tests on bean leaves two weeks after inoculation with sterile osmosed water, *Xcf* CFBP 13761 WT and *Xcf* CFBP 13761 *ΔhrcV*, respectively.**

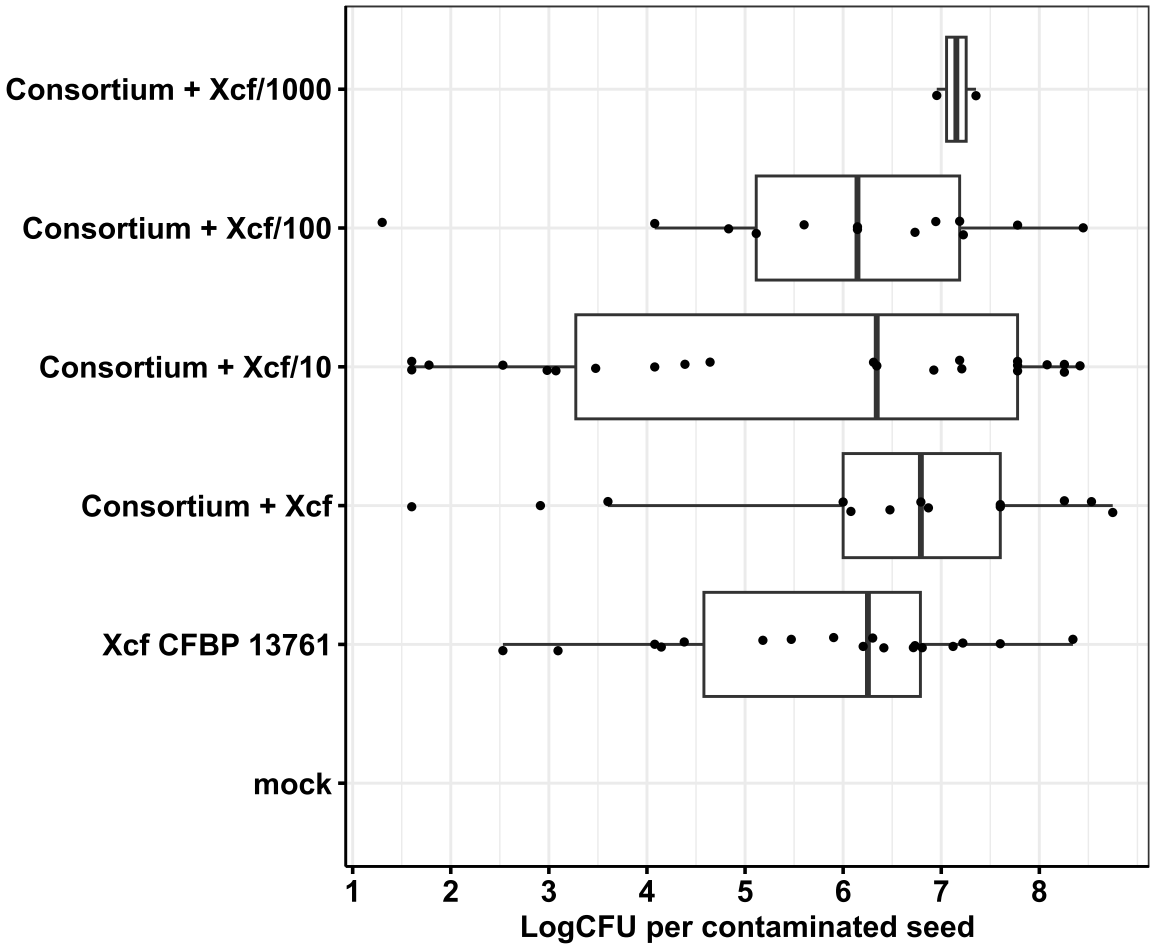


**Figure S4:** ***Xcf*** **population sizes on contaminated bean seeds inoculated with or without a ten bacterial strains consortium pure or diluted to a 1:10, 1:100 or 1:1000 ratio.**
